# Maternal heat stress induces persistent placental dysfunction and fetal growth restriction via the ERS-MAPK-apoptosis axis

**DOI:** 10.64898/2026.06.05.730306

**Authors:** Chang Yan, Chenjun Wang, Biyan He, Yifei Zhang, Siyu Wu, Yongtian Yin, Caixue Xu, Yiling Xiang, Yingjie Wu, Ning Liu, Yinghe Qin

## Abstract

Maternal heat stress (HS) is an emerging risk factor for adverse pregnancy outcomes, yet how gestational heat exposure causes persistent placental dysfunction remains unclear. In the present study, we established a murine HS model (38.5°C, 2.5 h/day, E0-E12.5) followed by thermal recovery to E17.5. HS reduced fetal weight during early gestation and caused persistent fetal growth restriction after recovery, despite partial placental weight restoration. Histological analyses revealed early reductions in the junctional and labyrinth zones, followed by sustained labyrinthine deficiency and compensatory junctional zone expansion. Consistently, HS impaired placental vascularization, with reduced vessel length and area, decreased CD31 and α-SMA abundance, and altered angiogenesis-related gene expression. HS also triggered oxidative stress, weakened antioxidant capacity, disrupted anti-inflammatory signaling, reduced tight junction protein expression, and compromised barrier integrity. Mechanistically, HS induced excessive endoplasmic reticulum stress, accompanied by increased CHOP, phosphorylated ERK, and cleaved caspase-3. In conclusion, our data unveil a heat-induced placental insufficiency program that restricts fetal growth through vascular, redox, barrier, and ERS-MAPK-apoptotic remodeling.

**Graphical Abstract:** 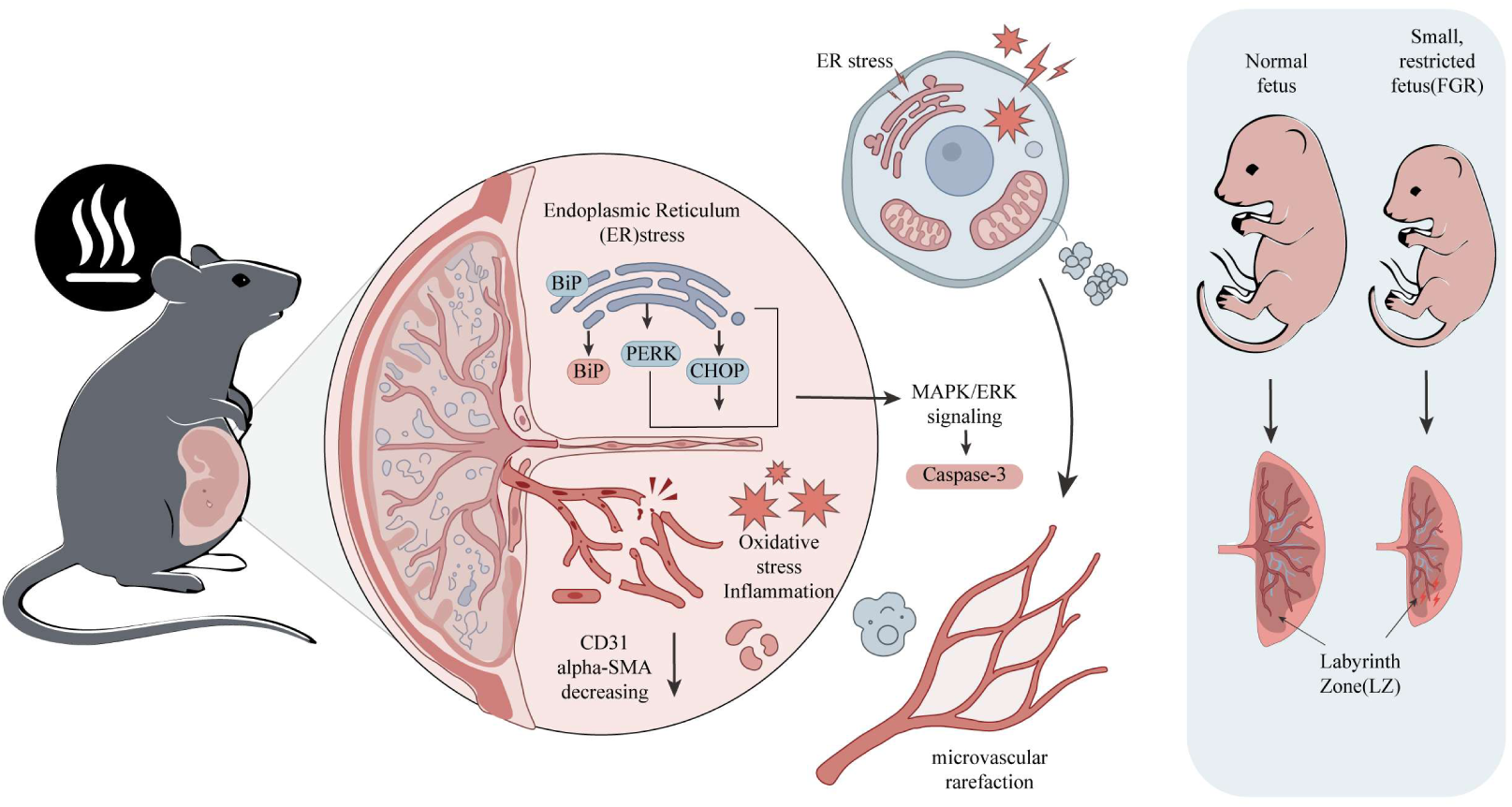

## Introduction

Global climate change has precipitated an unprecedented surge in the frequency and intensity of extreme heat events, posing a formidable threat to human reproductive health worldwide (Cramer et al, 2022; Ebi et al, 2021). Pregnant women exhibit a distinct vulnerability to hyperthermia due to altered thermoregulatory physiology and the metabolic demands of the growing fetus (Bonell et al, 2022; Jiao et al, 2023). Epidemiological evidence has consistently linked maternal heat exposure to adverse pregnancy outcomes, most notably fetal growth restriction (FGR) (Chersich et al, 2020; Essers et al, 2024; Ha et al, 2017). As a critical determinant of neonatal health, FGR predisposes offspring to immediate complications and long-term metabolic or cardiovascular sequelae (Barker 2004; Wu et al, 2006). However, translating these correlational findings into a clear mechanistic framework remains challenging. While human studies have demonstrated that each 2.4°C increase in ambient temperature raises the risk of low birth weight by approximately 18% (Ha et al, 2017), marked heterogeneity across populations is attributed to differences in climate adaptation and socioeconomic resilience (Ngo and Horton 2016; Sun et al, 2019). In contrast to the variability in human data, controlled animal models across multiple species provide a consistent demonstration that heat stress (HS) directly compromises fetal development (Monteiro et al, 2016). A pivotal observation in these models is that placental dysfunction often precedes fetal growth restriction (Bell et al, 1987). By contrast, subsequent studies have reported divergent placental responses to stress, including transient adaptive hypertrophy after stress relief (Galan et al, 1999) and irreversible structural defects (Zhao et al, 2020). This divergence points to a fundamental and unresolved question: whether the timing and duration of heat stress determine a threshold at which adaptive placental remodeling progresses to irreversible pathological damage. Emerging evidence points to the structural integrity of the placental vascular network as the decisive factor, where labyrinthine angiogenesis and mural cell recruitment dictate the perfusion capacity essential for fetal survival (Brosens et al, 2019; Hu et al, 2021; Hu et al, 2020; Ji et al, 2021; Ortega et al, 2022).

Mechanistically, maternal thermoregulation during HS involves peripheral vasodilation to facilitate heat dissipation, often at the functional expense of uteroplacental blood flow (Bonell et al, 2022). Chronic reductions in perfusion lead to impaired angiogenesis and malformed vascular architecture (Limesand et al, 2018). Paradoxically, while HS is linked to aberrant expression of key angiogenic regulators (e.g., Vegf, Plgf) and increased vascular resistance (Hafez et al, 2010; Olivier et al, 2021), some reports note increased vascular density alongside reduced cellularity, suggesting a maladaptive strategy to lower flow resistance (Nanas et al, 2021). These conflicting findings highlight a critical knowledge gap that the specific molecular mechanisms driving the shift from adaptive to maladaptive remodeling, as well as the exact time window for irreversible damage, have yet to be elucidated.

The function of placenta is largely determined by the meticulous development of the fetal microvascular network within the labyrinth zone, a process relying on the precise differentiation and anchoring of endothelial cells and pericytes (Rossant and Cross 2001). However, interrogation of signaling dynamics under HS uncovers a central biological paradox. While HS is well documented to trigger placental vascular rarefaction and pericyte loss (Armulik et al, 2011), stress-induced signaling cascades, such as the MAPK/ERK pathway, are frequently reported to be hyperactivated as part of the canonical cellular stress response (Kyriakis and Avruch 2012; Son et al, 2011).

In the present study, we hypothesize that this paradox is rooted in the pathological reprogramming of intracellular signaling driven by unresolved endoplasmic reticulum stress (ERS). We utilized a murine model of gestational HS and thermal recovery to delineate the temporal susceptibility of the placental vasculature. Our findings identify a novel ERS-MAPK-apoptosis signaling axis, where proteostatic exhaustion uncouples ERK signaling from its physiological role, transforming it into a driver of vascular dysfunction and barrier breakdown. By defining this critical developmental window, we offer a new molecular perspective on how environmental thermal extremes permanently program placental insufficiency and FGR.

## 2. Materials and methods

### 2.1 Ethics statement

All experiments involving animals were conducted according to the ethical policies and procedures approved by the China Agricultural University Animal Care and Animal Use Committee (AW40605202-1-02).

### 2.2 Animals

The 7-week-old C57BL/6J male and female mice were purchased from Spearhead Biotech Co., Ltd. (Beijing, China). The 7-week-old mice were housed at cages and maintained under standard environmental conditions of temperature (24±2°C), relative humidity (60%) underwent a 2-week adaptation stage before mating. The beginning of the gestation (embryonic 0.5 day, E 0.5 d) referred to female was confirmed by visualization of a vaginal plug.

### 2.3 Heat stress mouse model

The pregnant mice were divided into control (Con, n = 12) and HS (n = 12) groups. Mice in the Con group were maintained under 24±2°C. Mice in the HS groups were exposed to 38.5 °C from 12:00 to 14:30 using an artificial climate chest and returned to 24±2 °C for the remainder of the day.

### 2.4 H&E staining and placental zonation assessment

Paraffin-embedded mouse placentas were serially sectioned at a thickness of 5 μm. To evaluate placental layer architecture, sections were subjected to hematoxylin and eosin (H&E) staining using a commercial kit (C0105, Beyotime, China). Briefly, sections were deparaffinized in xylene (2 × 10 min) and rehydrated through a graded ethanol series (100%, 90%, 70%, and 50% for 5 min each). Tissues were stained with hematoxylin for 5 min, rinsed under running tap water for 15-20 min, and counterstained with eosin for 30 s. Subsequently, sections were dehydrated in an ascending ethanol series (70%, 80%, 90%, and 100% for 10 s each), cleared in xylene (2 × 3 min), and mounted with a resinous medium. The boundaries between the junctional zone (JZ) and the labyrinth zone (LZ) were manually delineated by two independent investigators and quantified with the Case Viewer software (3DHISTECH, Ltd, Hungary).

### 2.5 Immunofluorescence staining

Following deparaffinization and rehydration, tissue sections underwent heat-induced antigen retrieval by using citrate buffer (pH 6.0) for 20 min. Nonspecific binding was blocked by incubating sections with 5% bovine serum albumin for 2 h at room temperature. Sections were then incubated overnight at 4°C with a primary antibody against mouse CD31 (Cluster of Differentiation 31; endothelial cell marker; AF3628, R&D Systems, USA). After washing, sections were incubated for 1 h at room temperature with appropriate fluorophore-conjugated secondary antibodies. Nuclei were counterstained with 4′, 6-diamidino-2-phenylindole (DAPI; P0131, Beyotime, China) prior to mounting. To perform quantitative morphometric analysis, fluorescence images were acquired from three to five randomly selected non-overlapping fields per slide. Vascular parameters, specifically total vessel area and total network length, were computationally determined using AngioTool software, ensuring unbiased quantification across all experimental groups (Zudaire et al, 2011).

### 2.6 RNA sequencing and transcriptomic analysis

Total RNA was extracted and subjected to transcriptomic sequencing by Majorbio Bio-Pharm Technology (Shanghai, China). To preserve RNA integrity and biochemical stability, all samples were maintained and transported on dry ice. Differentially expressed genes (DEGs) were defined based on the thresholds of a two-tailed *P* value < 0.05 and a minimum |log_2_(fold change)| ≥ 1. Bioinformatics analysis, encompassing hierarchical clustering and Gene Ontology (GO) enrichment, was executed via the Majorbio Cloud Platform (https://cloud.majorbio.com). Comprehensive downstream analyses were further conducted to validate the reproducibility of mRNA expression profiles and elucidate their functional significance.

### 2.7 Quantitative RT-PCR

Total RNA was isolated using TRIzol Reagent (Sigma-Aldrich, USA) and reverse-transcribed into cDNA using the Fastking RT kit (Tiangen, China) following the manufacturer’s instructions. Quantitative PCR was performed on a CFX96 Touch Real-Time PCR Detection System (Bio-Rad, USA) using 2 × SuperReal PreMix Plus (SYBR Green, Tiangen). The thermal cycling profile consisted of an initial denaturation at 95°C for 15 min, followed by 40 cycles of 95°C for 10 s, 60°C for 20 s, and 72°C for 30 s. Primer efficiency was validated via standard curve analysis, and product specificity was confirmed by melting curve analysis. Relative gene expression was calculated using the 2^-ΔΔCt^ method, with GAPDH serving as the internal control. All primer sequences utilized in this study are detailed in the Supplementary Table S1.

### 2.8 Western blot

Immunoblotting was performed according to standard protocols. Briefly, placental tissues were lysed in RIPA buffer (Beyotime, Beijing, China) supplemented with 1% protease and phosphatase inhibitor cocktails. Protein concentrations were determined using a BCA Protein Assay Kit (Beyotime). Equivalent amounts of protein were resolved by SDS-PAGE and electrotransferred onto polyvinylidene fluoride membranes. After blocking with 5% non-fat milk, membranes were incubated with primary antibodies overnight at 4°C. Protein bands were visualized using enhanced chemiluminescence detection reagents and quantified by densitometric analysis using ImageJ software, with β-actin serving as the internal loading control. A comprehensive list of primary antibodies is provided in Supplementary Table S2.

### 2.9 Biochemical assays

Tissue samples were homogenized in ice-cold PBS and centrifuged at 3,000 × g for 30 min at 4°C. The resulting supernatants were collected for subsequent analysis. The levels of total antioxidant capacity (T-AOC), malondialdehyde (MDA), reduced glutathione (GSH), and oxidized glutathione (GSSG) were determined using commercial assay kits purchased from Beyotime Biotechnology (Shanghai, China). Specifically, the T-AOC kit, MDA kit, GSH assay Kit, and GSSG assay kit were used according to the manufacturer’s instructions. All analyses were performed strictly in accordance with the provided protocols.

### 2.10 Statistical analysis

Statistical analyses were performed using SPSS (version 27.0), and graphical representations were generated with GraphPad Prism (version 10.0). Sample sizes (n), representing the number of independent biological replicates, individual mice, or independent experimental repetitions, are explicitly stated in the respective figure legends. Data are presented as mean ± SEM. for parametric distributions or as median with 95% confidence intervals (CI) for non-parametric data. Statistical significance was defined as *P* < 0.05. Instances where 0.05 < *P* <0.1 were considered to indicate a potential numerical trend, albeit without reaching formal significance.

## 3. Results

### Maternal heat stress uncouples placental adaptation from fetal growth to drive persistent fetal growth restriction

To investigate the impact of climate-driven HS on mammalian reproduction, we established a murine model of thermal exposure (Figure 1A). Intriguingly, HS exposure did not significantly alter the live litter size at either embryonic day 12.5 (E12.5) or E17.5 (Figures 1B, C), suggesting that this specific thermal challenge protocol does not compromise overall fetal viability. However, average fetal weight was significantly diminished in the HS exposed group as early as E12.5 (Figures 1D, E). To assess placental development, we measured placental weight and calculated placental efficiency (fetal to placental weight ratio). At E12.5, HS significantly reduced absolute placental weight (Figure 1G), yet placental efficiency remained comparable to control levels (Figure 1H), indicating a proportional reduction in both fetal and placental mass during the initial stress phase.

**Figure 1.**
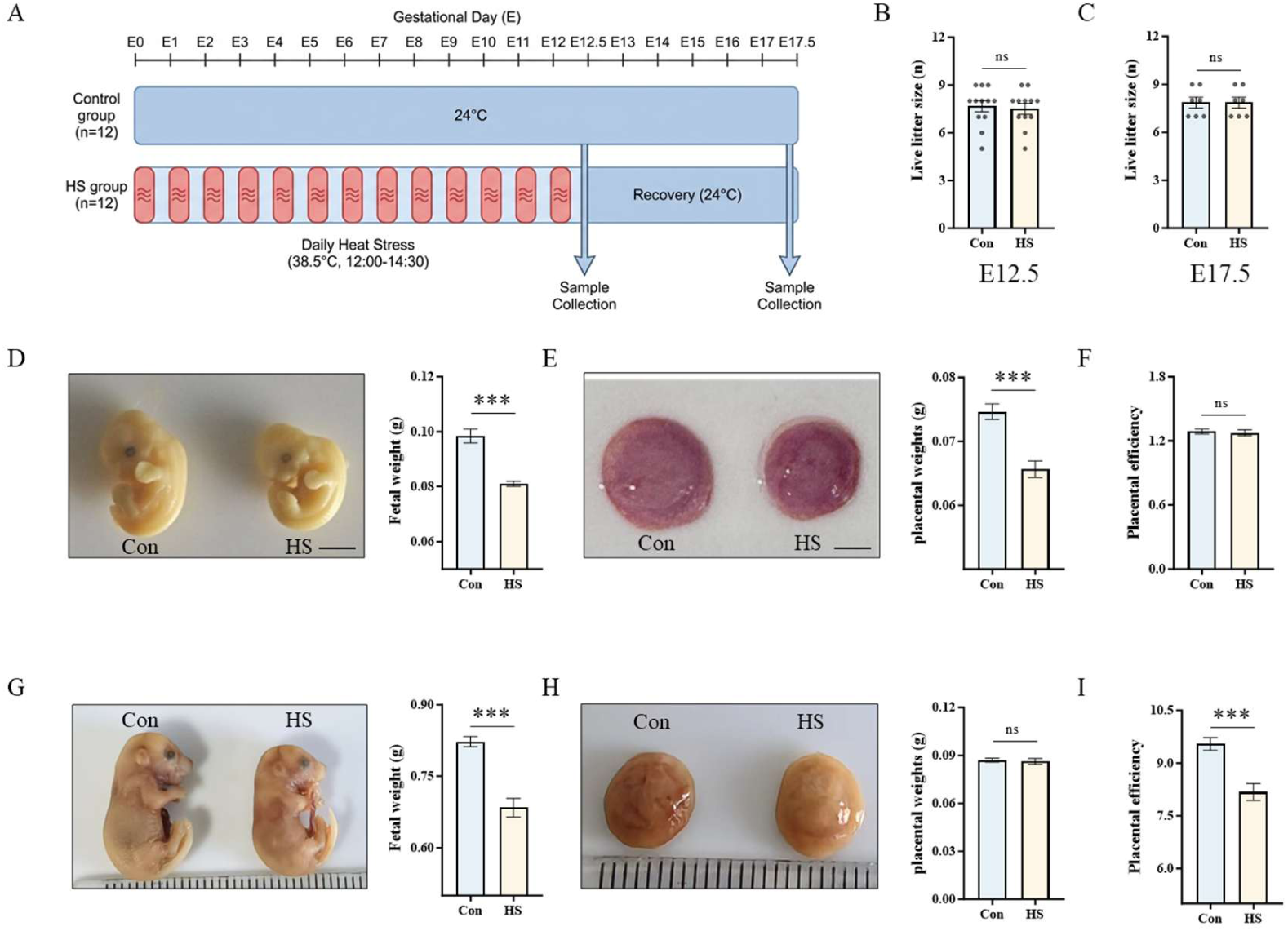
Heat stress induces persistent fetal growth restriction (FGR) and uncouples fetal-placental growth dynamics. (A) Schematic representation of the experimental design. Pregnant mice were subjected to either continuous heat stress (HS) until embryonic day 13 (E 12.5d) or an initial HS phase followed by a thermoneutral recovery period until E 17.5d. (B-C) Quantification of implantation sites and live litter size (B), and the calculated abortion rate (C) at E 12.5. (D) Representative images of fetuses (left) and quantification of average fetal weight (right) at E 12.5d. (E) Representative images of placentas (left), average placental weight (middle), and calculated placental efficiency (fetal weight/placental weight ratio, right) at E 12.5. (F) Quantification of live litter size at E 17.5 following the recovery period. (G-H) Representative morphological images and corresponding weights of fetuses (G) and placentas (H) at E 17.5d, alongside the quantitative assessment of placental efficiency (H, right panel). Data are presented as mean ± SEM. Statistical significance was determined using unpaired two-tailed Student’s t-test. ***, *P*< 0.001; ns, not significant. n = 12 dams per group.

To determine whether HS induced developmental impairment persists after the removal of the stressor, we subjected pregnant mice to HS until E12.5, followed by a thermoneutral recovery period until E17.5 (Figure 1A). Consistent with the early-gestational findings, the number of live fetuses remained stable through E17.5 (Figure 1C). Notably, while the absolute placental weight in the HS group rebounded and was no longer significantly different from controls by E17.5 (Figures 1K, L), fetal weight remained severely restricted (Figures 1I, J). Consequently, this uncoupling of placental recovery from fetal growth resulted in a sharp, statistically significant decline in placental efficiency at E17.5 (Figure 1M). These findings suggest that while the placenta retains a degree of gross structural plasticity enabling it to recover its mass once thermal stress is removed, its functional capacity remains irreversibly compromised. The stable placental efficiency observed at E12.5 likely reflects a transient state where fetal metabolic demands are still within the placenta’s reduced compensatory range. However, as gestation progresses into the rapid expansion phase, this underlying functional deficit is unmasked, failing to meet the escalating nutritional requirements of the fetus and ultimately culminating in profound FGR.

### Heat stress decouples labyrinth zone expansion from functional sinusoid development

To elucidate the morphological basis underlying the observed decline in placental efficiency and persistent FGR, we performed a detailed morphometric evaluation of the placental compartments. Histological analysis at E12.5 revealed that HS significantly altered the structural composition of the placenta: the area proportions of the JZ and the LZ were significantly reduced, whereas the proportion of the decidua was significantly increased (Figures 2A-D). These findings suggest that acute heat exposure during mid-pregnancy disproportionately suppresses the development of the fetal-derived compartments. By E17.5, the placental architecture exhibited a distinct structural shift. Although the proportion of the decidua returned to levels comparable with the control group, the LZ proportion remained significantly diminished in the HS group (Figure 2E-H). Conversely, the LZ proportion showed a compensatory increase compared to controls (Figure 2F). Crucially, the persistent reduction in the LZ proportion is particularly significant as the LZ is the primary site for maternal-fetal nutrient and gas exchange. The observation that HS leads to a contracted LZ, coupled with the previously noted reduction in maternal blood sinusoid area, strongly suggests that the functional deficit is rooted in impaired exchange interface development.

**Figure 2.**
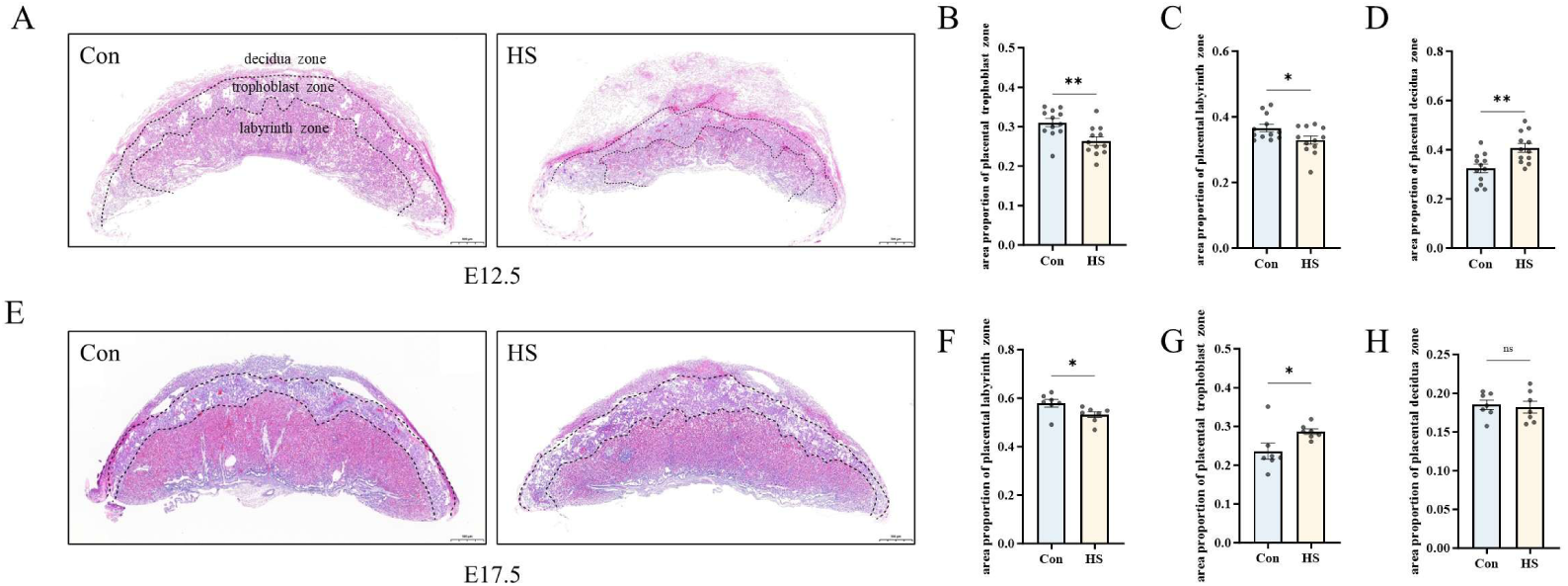
Heat stress induces dynamic remodeling of placental zonal proportions during mid-to-late gestation. (A, E) Representative images of H&E-stained placental sections at embryonic day 12.5 (E12.5; A) and E17.5 (E). Dashed lines delineate the boundaries of the decidua zone (DZ), junctional zone (JZ), and labyrinth zone (LZ). Scale bar = 500 um. (B-D) Quantitative morphometric analysis of placental compartments at E12.5. HS exposure significantly reduced the area proportions of the junctional zone (B) and labyrinth zone (C), while the proportion of the maternal decidua (D) was significantly increased. (F-H) Quantitative morphometric analysis of placental compartments at E17.5. At this stage, the area proportion of the labyrinth zone (F) in the HS group remained significantly lower than that of the control group. Conversely, the junctional zone proportion (G) exhibited a significant compensatory increase, while the decidua proportion (H) returned to control levels .Data are presented as mean ± SEM (n = 12 per group). Statistical significance was determined by unpaired Student’s t-test. \**P* < 0.05; \*\**P* < 0.01; ns, not significant.

### Heat stress impairs placental microvascular architecture and dysregulates angiogenic signaling

Given the persistent contraction of the labyrinthine compartment observed at E17.5, we hypothesized that the compromised transport capacity was rooted in a failure of the placental microvascular network. To characterize the cellular and molecular basis of this vascular impairment, we next evaluated endothelial and mural cell integrity. Immunofluorescence profiling of the endothelial marker CD31 demonstrated that HS significantly attenuated both vessel area and vessel length within the LZ at E12.5 (Figures 3A, C, D), a defect that persisted through E17.5 (Figure 3B), signifying a severe disruption of the endothelial network. These histological observations were corroborated by immunoblot analysis at E12.5, which confirmed a significant downregulation of CD31 protein expression (Figures 3E, G). Furthermore, the expression of the mural cell marker α-SMA was also significantly reduced in HS placentas (Figures 3E, H), indicating that the structural stability provided by pericytes or smooth muscle cells was compromised. To determine whether this profound vascular breakdown was orchestrated at the transcriptional level, we assessed the mRNA expression of key vascular regulators via RT-qPCR at E12.5 (Figures 3F). Consistent with our protein-level findings, HS induced a significant downregulation of the smooth muscle marker *Cnn1* and the critical remodeling factor *Fgfr1* (Figures 3F), further validating the extensive depletion of the mural compartment. Notably, we uncovered a striking decoupling within canonical angiogenic signaling pathways. While the pro-angiogenic ligand *Plgf* was markedly upregulated in the HS group, its cognate receptors (*Flt1*, *Vegfr2*) failed to mount a compensatory response, remaining unchanged between groups (Figure 3F). Collectively, these data establish that HS drives a comprehensive breakdown of the placental vascular architecture by compromising both endothelial and mural compartments. This structural failure appears to be driven by a mismatch in angiogenic signaling, where a paradoxical surge in *Plgf* is insufficient to rescue the impaired vascular remodeling and maintenance.

**Figure 3.**
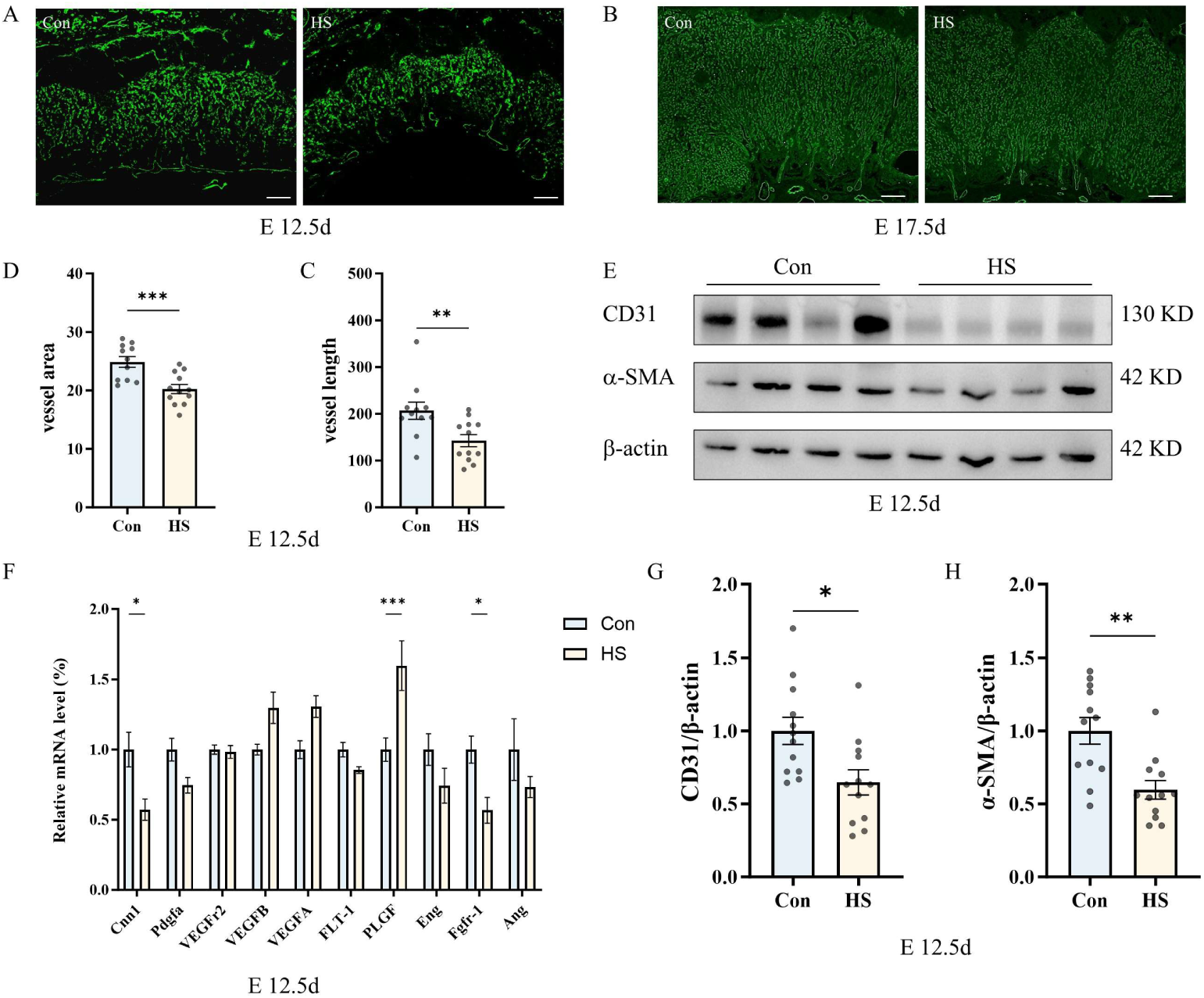
Heat stress impairs placental vascularization and dysregulates angiogenic signaling. (A, B) Representative immunofluorescence images of CD31 (green) in the placental labyrinth zone (LZ) at E12.5 (A) and E17.5 (B). Scale bars = 200 um. (C, D) Morphometric quantification of vessel length (C) and vessel area (D) at E12.5, showing a significant reduction in vascular density in the HS group. (E) Western blot analysis of endothelial marker CD31 and mural cell marker α-SMA in E12.5 placental tissues. β-Actin served as the loading control. (G, H) Densitometric quantification of CD31 (G) and α-SMA (H) protein levels normalized to β-Actin (n = 12 per group). (F) Relative mRNA expression levels of vascular and angiogenic-related genes (*Cnn1*, *Pdgfa*, *Vegfr2*, *Vegfb*, *Vegfa*, *Flt-1*, *Plgf*, *Eng*, *Fgfr1*, and *Ang*) at E12.5 determined by RT-qPCR. Expression levels were normalized to GAPDH. Data are presented as Mean ± SEM. Statistical significance was determined by unpaired Student’s t-test; \**P* < 0.05, \*\**P* < 0.01, \*\*\**P* < 0.001.

### Heat stress induced oxidative damage and inflammation response dysregulation compromise placental barrier integrity

Considering that the placental vasculature is highly sensitive to the local biochemical milieu, we reasoned that the observed vascular collapse might be secondary to a deteriorating placental microenvironment. To delineate the upstream drivers of this failure, we evaluated markers of oxidative stress, inflammatory signaling, and barrier integrity. HS induced a profound state of oxidative stress, characterized by a significant accumulation of the MDA (Figure 4A). This was accompanied by a comprehensive collapse of the antioxidant defense system, evidenced by the significant depletion of T-AOC, SOD activity, and the GSH/GSSG ratio (Figures 4B-E). Concurrently, HS precipitated a robustly polarized immunological response. RT-qPCR revealed a marked induction of the pro-inflammatory cytokines *Tnf-α* and *Il-1β*, occurring alongside a profound suppression of the anti-inflammatory mediators *Tgf-β* and *Il-10* (Figure 4F). Notably, *IL-6* expression was also significantly downregulated. This atypical cytokine profile suggests an uncoupling of systemic immune coordination, characterized by a specific deficit in *IL-6* mediated signaling. We hypothesized that this skewed, pro-oxidant, and inflammatory microenvironment directly compromises the structural integrity of the placental barrier. To investigate this, we assessed the expression of key tight junction proteins. Immunoblot analysis demonstrated a significant downregulation of ZO-1, Claudin-11, and Occludin in HS exposed placentas (Figures 4G-J). Collectively, these findings indicate that HS-induced oxidative damage and immunological dysregulation orchestrate the breakdown of the maternal-fetal barrier, creating a hostile environment that exacerbates vascular impairment and ultimately culminates in fetal growth restriction.

**Figure 4.**
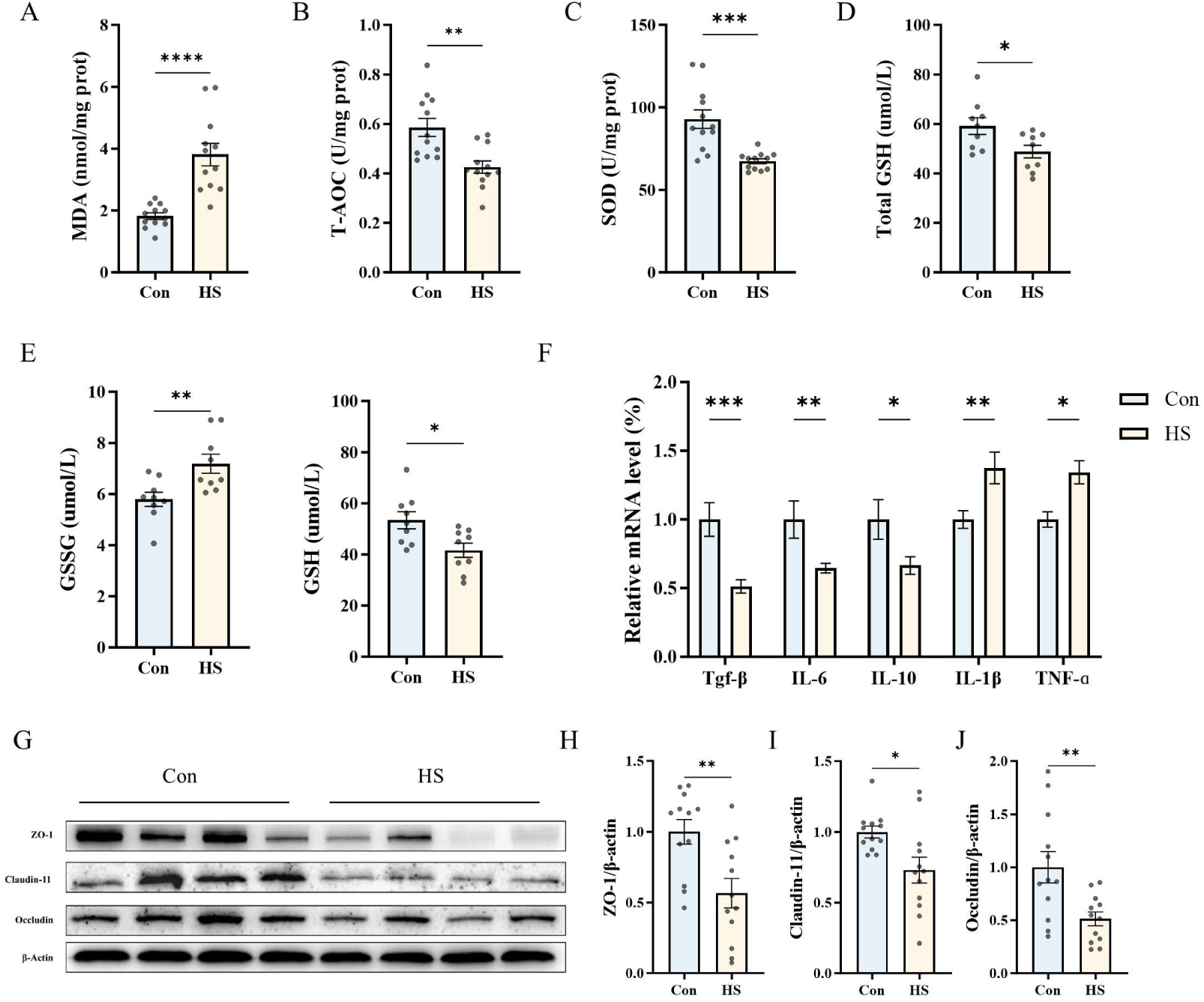
Heat stress triggers oxidative stress, inflammatory dysregulation, and compromises placental barrier integrity. (A-E) Biochemical assessment of placental oxidative stress markers. (A) Levels of malondialdehyde (MDA), a marker of lipid peroxidation. (B-D) Evaluation of the antioxidant defense system, including total antioxidant capacity (T-AOC, B), superoxide dismutase (SOD) activity (C), and total glutathione (GSH) levels (D). (E) Concentrations of oxidized glutathione (GSSG) and reduced glutathione (GSH). (F). Relative mRNA expression levels of anti-inflammatory mediators in placental tissues, normalized to GAPDH. (G-J) Protein expression of placental tight junction markers. (G) Representative Western blot images for ZO-1, Claudin-11, and Occludin, with β-Actin as the loading control. (H-J) Densitometric quantification of ZO-1 (H), Claudin-11 (I), and Occludin (J) protein levels normalized to β-Actin. Data are presented as Mean ± SEM (n = 12 per group). Statistical significance was determined by unpaired Student’s t-test; \**P* < 0.05, \*\**P* < 0.01, \*\*\**P* < 0.001.

### Heat stress drives placental vascular failure through an integrated ER stress-centered transcriptional program

To identify the molecular events underlying HS-induced placental vascular failure, we performed KEGG enrichment analysis of placental DEGs. The total DEG set was enriched for pathways involved in xenobiotic metabolism, cytochrome P450-related drug metabolism, arginine biosynthesis, glutathione metabolism, cAMP signaling, Hippo signaling and fatty acid degradation (Fig. 5A), indicating broad disruption of vascular tone, endothelial redox control, barrier function and metabolic support. Direction-specific enrichment analysis revealed a clear separation between stress-activated and homeostatic pathways. Upregulated genes were enriched in protein processing in the endoplasmic reticulum, MAPK signaling, endocytosis, cytokine-cytokine receptor interaction, JAK-STAT signaling and PI3K-Akt signaling (Fig. 5B), consistent with activation of a proteotoxic, inflammatory and maladaptive angiogenic stress response. Conversely, downregulated genes were enriched in cytochrome P450 metabolism, arginine biosynthesis, glutathione metabolism, cAMP signaling, Hippo signaling and fatty acid degradation (Fig. 5C), pathways required for endothelial detoxification, nitric oxide-dependent vascular tone, antioxidant defense and metabolic homeostasis. The chord plot further showed convergence of DEGs on ER protein processing, MAPK, PI3K-Akt, cytokine and metabolic pathways, positioning ER stress as a central node connecting thermal injury to placental vascular dysfunction (Fig. 5D).

**Figure 5.**
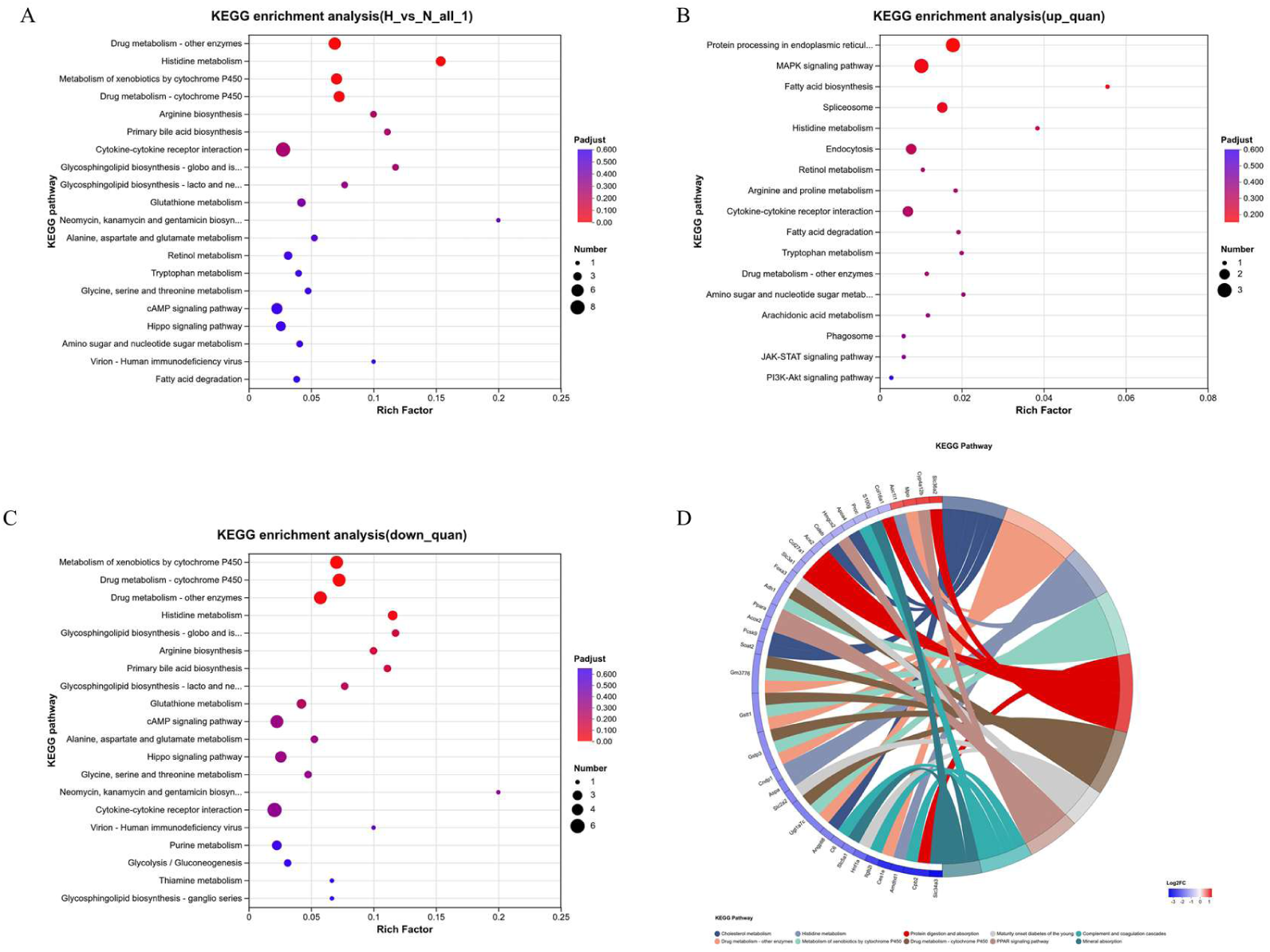
Transcriptomic profiling of heat stress-exposed placentas identifies ER stress and metabolic disruption as central pathological features. (A) KEGG pathway enrichment analysis of all differentially expressed genes (DEGs) in heat stress (HS)-treated placentas compared to control (Con) tissues. Pathways are ranked by their rich factor; dot size represents the number of genes enriched in each pathway, and color indicates the significance level (Padjust). (B, C) Stratified KEGG enrichment analysis showing the top functional pathways for upregulated DEGs (B) and downregulated DEGs (C). Upregulated pathways (B) highlight the activation of protein processing in the endoplasmic reticulum (ER), MAPK signaling, and inflammatory cascades. Downregulated pathways (C) reveal suppressed metabolic programs, including arginine biosynthesis, glutathione metabolism, and vascular-regulatory cAMP/Hippo signaling. (D) KEGG chord plot illustrating the relationship between specific DEGs and key functional pathways (e.g., ER protein processing, MAPK signaling, and metabolic pathways). The outer ring shows the gene symbols, colored by their log_2_2 (fold change) (blue for downregulation, red for upregulation). Inner ribbons connect genes to their respective KEGG pathway categories, highlighting the interconnected nature of the HS-induced stress response.

### Heat stress drives placental failure through an ERS-MAPK-apoptosis axis

To substantiate the transcriptomic signatures, we quantified the expression of key regulators governing placental vascular homeostasis and metabolic transport via qPCR. Our results confirmed that HS exposure selectively compromised the placental vascular-metabolic unit rather than inducing a generalized transcriptional collapse. While the expression of *Angptl8* a regulator involved in angiogenesis and lipid partitioning remained stable (Fig. 6A), HS significantly suppressed the transcripts of *Ace2*, a pivotal component for vascular protection and hemodynamic stability (Fig. 6B). This vascular impairment was coupled with a profound disruption of metabolic programs. Specifically, we observed the marked downregulation of *Ppara*, *Hmgcs2*, and *Apoa4* (Fig. 6C-E), indicating a systemic failure of lipid metabolic pathways and transport mechanisms essential for endothelial energy homeostasis. Furthermore, the expression of the detoxification enzyme *Ugt1a7c* was significantly reduced (Fig. 6F), while *Adh1* exhibited a downward trend (Fig. 6G), reflecting a compromised capacity for placental metabolic processing. Conversely, *Cyp4a12b* was significantly induced under HS (Fig. 6H), potentially signifying a pathological shift toward vasoconstrictive arachidonic acid metabolism. Collectively, these data suggest that HS selectively silences the molecular machinery required for vascular maintenance and metabolic exchange while promoting maladaptive stress responses.

**Figure 6.**
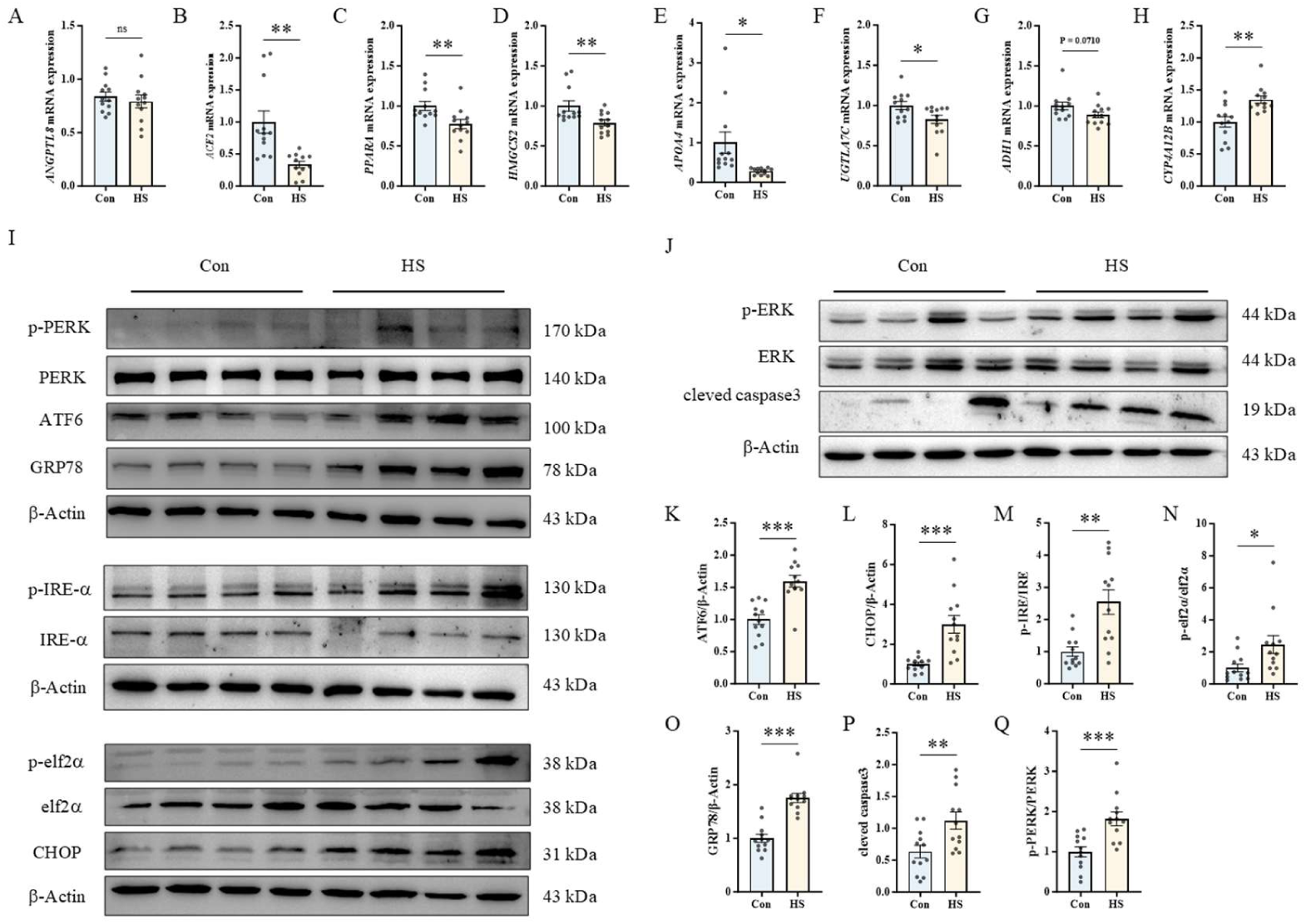
Activation of the ERS-MAPK-apoptosis axis mediates heat stress-induced placental vascular and metabolic injury. (A-H) Quantitative PCR (qPCR) validation of transcriptomic signatures. Relative mRNA expression levels of genes governing angiogenesis (*ANGPTL8*, A), vascular protection (*ACE2*, B), lipid metabolic homeostasis (*PPARA*, C; *HMGCS2*, D; *APOA4*, E), metabolic detoxification (*UGT1A7C*, F; *ADH1*, G), and pathological arachidonic acid metabolism (*CYP4A12B*, H) in control (Con) and heat-stressed (HS) placentas. (I, J) Representative Western blot images showing the activation of the endoplasmic reticulum (ER) stress machinery and downstream signaling cascades. Blots display key components of the unfolded protein response (UPR) (I) alongside markers for MAPK signaling (ERK) and terminal apoptosis (cleaved caspase-3) (J). β-Actin served as the loading control. (K-Q) Densitometric quantification of the immunoblots. The data confirm the robust induction of the ER chaperone GRP78 (O) and the activation of all three UPR branches: increased ATF6 (K), phosphorylated IRE1α (M), phosphorylated PERK (Q), and its downstream target phosphorylated eIF2α (N). This unresolved proteotoxic stress culminates in the upregulation of the pro-apoptotic transcription factor CHOP (L) and the apoptotic executioner cleaved caspase-3 (P). Data are presented as mean ± SEM, n=12 biologically independent samples per group. Statistical significance was determined by two-tailed unpaired Student’s t-test. \**P* < 0.05, \*\**P* < 0.01, \*\*\**P* < 0.001; ns, not significant.

Guided by the prominent enrichment of proteostasis-related pathways, we next examined the activation of the ERS machinery at the protein level. HS placentas exhibited a robust ERS signature, characterized by a significant induction of the molecular chaperone GRP78 (BiP), a definitive marker of severe proteotoxic stress (Fig. 6I, K). This was accompanied by the potent activation of all three canonical branches of the unfolded protein response (UPR): increased ATF6 abundance and enhanced phosphorylation of IRE1α and PERK, as well as its downstream target eIF2α (Fig. 6I, L-N). The transition from adaptive signaling to terminal cell death was evidenced by the marked upregulation of the pro-apoptotic transcription factor CHOP (Fig. 6I, O), suggesting that the unresolved proteostatic burden under HS precipitates irreversible tissue injury.

We further explored the propagation of this stress to downstream signaling cascades. Paradoxically, despite the overt vascular rarefaction, HS placentas exhibited an aberrant hyperactivation of ERK signaling, as evidenced by significantly increased p-ERK/ERK ratios (Fig. 6J, P). This uncoupling of robust ERK phosphorylation from functional vascularization suggests that HS pathologically reprograms the MAPK pathway, potentially shifting it from a pro-survival angiogenic signal to a driver of cellular dysfunction. This maladaptive molecular environment culminated in the activation of the apoptotic executioner, cleaved caspase-3 (Fig. 6J, Q). Together, these findings establish that HS drives placental failure through an ERS-MAPK-apoptosis axis, in which unresolved UPR signaling decommissions vascular-metabolic homeostasis and triggers widespread programmed cell death, ultimately leading to vascular rarefaction and impaired placental perfusion.

## Discussion

Maternal HS is emerging as a major environmental determinant of adverse pregnancy outcomes in the context of global warming (Cramer et al, 2022; Ebi et al, 2021), yet the biological basis by which transient thermal insult is converted into sustained placental dysfunction remains incompletely understood. In this study, we demonstrate that gestational HS does not primarily impair pregnancy maintenance, but instead induces a persistent placental failure phenotype that culminates in FGR. By integrating gross phenotyping, placental morphometry, vascular profiling, redox and barrier analyses, transcriptomics, and protein-level validation, we identify a mechanistic cascade in which HS progressively disrupts placental structural organization, compromises labyrinthine vascularization, destabilizes maternal-fetal barrier integrity, and activates a terminal ERS-MAPK-apoptosis program. These findings support a model in which the placenta initially undergoes an apparently compensatory but ultimately maladaptive response to heat exposure, resulting in permanent functional insufficiency even after restoration of a thermoneutral environment.

A key finding of the present study is that HS uncouples placental growth dynamics from fetal growth outcome. Although maternal HS significantly reduced fetal weight at E 12.5 d, it did not alter implantation number, live litter size, or abortion rate, indicating that the thermal paradigm used here was sufficient to impair fetal development without causing overt embryonic loss. This is consistent with previous research reports (Zheng et al, 2023). At this stage, placental weight was also reduced, whereas placental efficiency remained relatively stable, suggesting that early HS suppresses fetal and placental growth in parallel. However, this apparent proportionality was lost after thermal recovery. By E17.5 d, placental weight had substantially rebounded, yet fetal weight remained markedly reduced, producing a pronounced decline in placental efficiency. This divergence is conceptually important because it indicates that restoration of placental mass does not equate to restoration of placental competence. Similar dissociations between placental size and functional capacity have been noted in other forms of placental stress, where gross tissue recovery masks persistent defects in exchange structure, vascular function, or cellular differentiation (Higgins et al, 2016; Woods et al, 2018). Our data extend this concept to gestational HS and indicate that placental adaptation to heat is not merely incomplete, but fundamentally misaligned with fetal demand.

The morphometric analyses further clarify the structural basis of this functional uncoupling. At E12.5, HS reduced both the JZ and LZ while relatively expanding the decidual compartment, indicating that heat exposure preferentially suppresses development of fetal-derived placental regions. The reduction in LZ is of particular importance, as this compartment contains the specialized maternal-fetal exchange interface required for nutrient, gas, and waste transport (Woods et al, 2018). By E17.5, the decidual proportion had normalized and the JZ displayed an apparent compensatory increase, yet the LZ deficit persisted. This spatially selective remodeling suggests that the placenta retains limited plasticity after removal of the heat insult, but that such compensation does not reconstruct the exchange architecture most essential for fetal growth. In this regard, our findings help reconcile previous discrepancies in the literature, where some studies have reported placental hypertrophy or partial recovery after stress withdrawal, whereas others have emphasized irreversible structural injury (Morrison 2008; Sferruzzi-Perri et al, 2017). The present data suggest that both observations may be correct, but reflect different levels of analysis: bulk placental mass may partially recover, whereas the labyrinthine microstructure, once perturbed during a sensitive developmental period, may fail to regenerate adequately.

The vascular analyses identify labyrinthine angiogenic failure as a central lesion underlying persistent placental insufficiency. HS markedly reduced vessel length and vessel area within the LZ and downregulated CD31 protein expression, indicating significant endothelial loss and microvascular rarefaction. At the same time, α-SMA expression was decreased, implying impaired mural cell recruitment or stabilization of nascent vessels. Because functional placental angiogenesis requires coordinated interactions among endothelial cells, perivascular support cells, trophoblasts, and angiogenic ligands, these findings indicate a broader collapse of vascular maturation rather than a simple delay in vessel expansion (Cross et al, 2006). This interpretation is reinforced by the dysregulated expression of angiogenesis-associated genes, including mediators linked to VEGF signaling, vascular remodeling, and vessel stabilization. Prior studies have shown that maternal hyperthermia or reduced uteroplacental perfusion can disrupt placental vascular development and increase fetal vulnerability (Burton and Jauniaux 2018). Our results advance this framework by demonstrating that HS simultaneously impairs both endothelial abundance and mural support, thereby compromising not only vascular density but also vessel integrity. This may explain why placental tissue can regain mass without regaining effective exchange performance: the limiting defect resides in the organization and maturity of the vascularized exchange interface, not simply in placental size.

The oxidative stress data provide a compelling upstream context for these structural defects. HS increased placental MDA while decreasing T-AOC, SOD activity, and glutathione buffering capacity, indicating severe redox disequilibrium. Given the exceptionally high metabolic demands of the placenta, sustained oxidative burden is likely to be particularly deleterious in this tissue, where redox imbalance can disrupt membrane lipids, mitochondrial function, and signaling networks essential for vascular and trophoblast homeostasis (Aouache et al, 2018). Oxidative stress has been implicated in a wide range of placental pathologies, including FGR, preeclampsia, and ischemia-related injury (Burton and Jauniaux 2011). The current findings place oxidative injury prominently within the heat stress response and support the view that excessive reactive oxygen species generation is not simply a byproduct of placental dysfunction, but a potential upstream driver of maladaptive remodeling. This is mechanistically relevant because oxidative stress is a potent inducer of ER dysfunction, protein misfolding, and stress kinase activation, thereby providing a plausible bridge between external thermal exposure and intracellular signaling collapse (Malhotra and Kaufman 2007).

In parallel with redox imbalance, HS altered the placental inflammatory milieu and weakened barrier-associated tight junction architecture. Although the precise directionality and cellular sources of these immune changes require further resolution, the altered expression of anti-inflammatory mediators suggests a loss of immunoregulatory equilibrium rather than maintenance of a protective anti-inflammatory state. This is likely to be biologically meaningful, as placental vascular remodeling, trophoblast differentiation, and barrier function are highly sensitive to local inflammatory context (Aplin 2010). Consistent with this, HS significantly reduced the expression of ZO-1, Claudin-11, and Occludin, indicating that the structural integrity of the maternal-fetal barrier was compromised. Tight junction disruption in the placenta has been linked to increased permeability, impaired selective transport, and heightened vulnerability to inflammatory and oxidative insults (Adu-Gyamfi et al, 2021). In the setting of HS, these barrier abnormalities likely synergize with vascular rarefaction to destabilize the exchange microenvironment. Thus, oxidative stress, inflammatory disequilibrium, and barrier failure should not be viewed as isolated events, but as interdependent elements of a progressive placental injury program.

The transcriptomic and biochemical data identify unresolved ERS as the central molecular node integrating these upstream perturbations. KEGG enrichment analysis highlighted pathways related to protein processing in the endoplasmic reticulum and MAPK signaling, implying that proteostatic disruption is a major feature of the heat-stressed placenta. This conclusion was strongly substantiated by immunoblotting, which showed elevated GRP78, activation of ATF6, phosphorylation of IRE1α, PERK, and eIF2α, and robust induction of CHOP. Collectively, these changes indicate activation of all three canonical UPR branches, culminating in terminal pro-apoptotic ERS (Tabas and Ron 2011). This distinction is critical. Under physiological or transient stress conditions, UPR activation serves an adaptive function by reducing translational load, increasing folding capacity, and restoring ER homeostasis. However, when the stress burden is excessive or prolonged, the UPR shifts from a pro-survival program to a pro-death program mediated in part by CHOP, mitochondrial dysfunction, and caspase activation (Tabas and Ron 2011). Our data strongly favor the latter scenario, indicating that HS drives the placenta beyond adaptive proteostasis and into a state of proteostatic exhaustion.

One of the most important conceptual advances of this work is the identification of pathological MAPK reprogramming within this terminal ERS context. ERK signaling is classically associated with cell proliferation, differentiation, and angiogenic adaptation, and in the developing placenta it is generally considered supportive of growth and survival (Kalisch-Smith 2025). However, we found that HS increased ERK phosphorylation in parallel with CHOP induction and cleaved caspase-3 accumulation, despite clear evidence of vascular rarefaction, barrier disruption, and persistent fetal growth impairment. These observations suggest that ERK activation under severe heat-induced stress is uncoupled from its canonical trophic function. Rather than mediating successful adaptation, ERK appears to be repurposed into a maladaptive signaling state that coexists with, and possibly amplifies, terminal ERS and apoptosis. This context-dependent reinterpretation of ERK signaling may help explain why activation of canonical “pro-growth” pathways does not necessarily predict improved placental outcome under pathological conditions (Wang et al, 2025). More broadly, our findings support the notion that the biological meaning of pathway activation cannot be inferred in isolation; instead, it must be understood within the proteostatic and redox landscape in which it occurs.

## Conclusion

In summary, the present study demonstrates that maternal HS induces persistent FGR not by simply reducing placental growth, but by triggering irreversible structural and molecular reprogramming of the placenta. HS causes sustained contraction of the labyrinthine exchange compartment, endothelial and mural cell loss, oxidative and inflammatory imbalance, barrier dysfunction, and activation of a terminal ERS-MAPK-apoptosis axis. These events uncouple placental recovery in mass from recovery in function and define a vulnerable developmental window beyond which damage becomes refractory to thermal normalization. By establishing this mechanistic framework, our work provides a stronger biological basis for understanding heat-associated placental insufficiency and suggests that preserving proteostasis, vascular integrity, and barrier function during early gestation may represent a promising strategy for mitigating the reproductive consequences of global warming.

## Abbreviations

BiP: GRP78
CD31: cluster of differentiation 31
E12.5: embryonic day 12.5
E17.5: embryonic day 17.5
ERS: endoplasmic reticulum stress
FGR: fetal growth restriction
GSH: glutathione
GSSG: oxidized glutathione
HS: heat stress
LZ: labyrinth zone
JZ: junctional zone
MDA: lipid peroxidation product malondialdehyde
SOD: superoxide dismutase
T-AOC: total antioxidant capacity
UPR: unfolded protein response.

## Author contributions

All the authors contributed to the writing of this review.

## Data availability

RNA-seq data generated for this study have been deposited in the NCBI Sequence Read Archive (SRA) with the accession number SUB16227148.

## Acknowledgements

This work was supported by the National Key Research and Development Program of China (2023YFD1302005), National Natural Science Foundation of China (No.32000082), R&D Program of Beijing Municipal Education Commission (KM20221244800), Earmarked Fund for Modern Agro-Industry Technology Research System (CARS-43), China Postdoctoral Science Foundation (No.2022M713405), and the 2115 Talent Program of China Agricultural University.

